# Spatiotemporal Dynamics of the Bacterial Microbiota on Lacustrine *Cladophora Glomerata* (Chlorophyta)

**DOI:** 10.1101/096347

**Authors:** Michael J. Braus, Linda E. Graham, Thea L. Whitman

## Abstract

The branched periphytic green alga *Cladophora glomerata,* often abundant in nearshore waters of lakes and rivers worldwide, plays important ecosystem roles, some mediated by epibiotic microbiota that benefit from host-provided surface, organic C, and O_2_. Previous microscopy and high throughput sequencing studies have indicated surprising epibiont taxonomic and functional diversity, but have not included adequate consideration of sample replication or the potential for spatial and temporal variation. Here we report the results of 16S rRNA amplicon-based phylum-to-genus taxonomic analysis of *Cladophora*-associated bacterial epibiota sampled in replicate from three microsites and at six times during the open-water season of 2014, from the same lake locale (Picnic Point, Lake Mendota, Dane Co., WI, USA) explored by high throughput sequencing studies in two previous years. Statistical methods were used to test null hypotheses that the bacterial community: 1) is homogeneous across microsites tested, and 2) does not change over the course of a growth season or among successive years. Results indicated a dynamic microbial community that is more strongly influenced by sampling day during the growth season than by microsite variation. A surprising diversity of bacterial genera known to be associated with the key function of methane-oxidation (methanotrophy)-including relatively high-abundance of *Crenothrix, Methylomonas,* and *Methylocaldum*–showed intra-seasonal and inter-annual variability possibly related to temperature differences, and microsite preferences possibly related to variation in methane abundance. By contrast, a core assemblage of bacterial genera seems to persist over a growth season and from year-to-year, possibly transmitted by a persistent attached host resting stage.

## Introduction

The periphytic filamentous green alga *Cladophora* commonly occurs in marine and freshwaters worldwide, is particularly abundant in environments affected by eutrophication, and plays important roles in ecosystem materials cycling and providing habitat for other organisms (reviewed by Zulkifly et al. 2013). Some of these ecosystem roles are mediated by epibiotic microorganisms, which benefit from high host surface area for attachment and host-generated organic carbon and oxygen. A previous high throughput 16S rRNA gene amplicon analysis indicated that the bacterial microbiota occupying surfaces of lacustrine *Cladophora glomerata* displayed high levels of taxonomic and inferred functional diversity, including cyanobacteria and other bacterial genera, many of which are known to be associated with methane-oxidation, nitrogen-fixation, and cellulose-degradation (Zulkifly et al. 2012). A subsequent metagenomic analysis of *Cladophora* sampled from the same locale revealed high archaeal, bacterial, and eukaryotic epibiont taxonomic diversity, as well as diverse functional genes such as prokaryotic *nifH* (a marker for nitrogen-fixation), *amo* (ammonia monooxygenases), genes involved in aerobic and anaerobic biosynthesis of vitamin B_12_ required by the host, and prokaryotic and eukaryotic genes encoding putative cellulases (Graham et al. 2015a). These results indicated that the *Cladophora* host and attached microbiota modify and exchange key materials relevant to N and C cycling.

Even so, Graham et al. (2015b) pointed out the need for greater attention to spatial and temporal replication of high throughput genetic sequencing studies of algal microbiota and microbiomes. These authors also recommended that greater attention be paid to the possibility that epibiotic microbiota might provide functional redundancy that could support holobiont response to environmental change. To address these issues we used 16S rRNA gene amplicon sequencing to classify and compare bacterial microbiota of *Cladophora glomerata* sampled in triplicate, and at six time points during the growth season of 2014, from three microsites at the same lacustrine locale employed in two previous high-throughput (but less-well replicated) studies (Zulkifly et al. 2012, Graham et al. 2015a). We used statistical methods to test the null hypotheses that the *C. glomerata* epibiotic bacterial community: 1) was relatively homogeneous across microsites tested, and 2) did not change substantially over the course of a growth season. We aimed to determine whether seasonality or small-scale geographical variation was the more important driver of epibiotic bacterial community variation during a growth season. We also used class‑ and genus-level comparisons of taxonomic data for the same locale from three years (2011, 2012, 2014) to explore inter-annual variability. Additionally, we used comparisons at the genus level of *Cladophora*-associated methanotrophic bacteria among these three years to gain insight into community environmental responsiveness and functional redundancy.

## MATERIALS AND METHODS

### Sampling locale and microsites

The sampling locale was Picnic Point, a shoreline peninsular feature of Lake Mendota, Dane Co., WI (43°4.53' N, W 89°35.5' W, Supplementary Figure 1). This locale was chosen because two previous high throughput studies of *Cladophora* microbiota (Zulkifly et al. 2013, Graham et al. 2015a) were based on samples obtained at the same place in 2011 and 2012, potentially allowing inter-annual comparisons. The locale was suitable for all three studies because it is relatively undisturbed by shoreline development, and is representative of other relatively undisturbed shorelines of the same lake (Graham et al. 2015a). Attached biomass dominated by the species *C. glomerata* (Graham et al. 2015a) was sampled six times between Julian days 172-214 (June 21^st^ to August 2^nd^), 2014 from three peninsular microsites, a northern aspect (N), southern aspect (S), and point (P), ∼100m apart. The point microsite was the least shaded by shoreline vegetation. Although *Cladophora* occurs on substrata located near the water surface as well as substrata located more deeply in the water column, for this study, sampling was restricted to depths of 1.0–1.5m. Water temperature at the depth of algal collection increased over the open-water period; water temperature was 21°C on Julian day 160 and peaked at 26°C on Julian day 220 (Supplementary Figure 2). Near-shore regions of Picnic Point and other areas of Lake Mendota examined in 2014 were supersaturated with methane; methane concentrations varied from 190 ppm at mid-summer, peaked at 350 ppm in late August, then dropped to 20 ppm in October (Luke Loken, University of Wisconsin Center for Limnology, personal communication).

### Sample collection, transport and processing

Using ethanol-sterilized tools, we removed three entire highly-branched thalli, including holdfasts, from submersed substrata and placed them in sterile Whirl-Pak bags (Nasco, Fort Atkinson, WI) that also contained local lake water and a headspace of air. Samples were transported to the lab in an insulated container, then processed the same day. To remove planktonic and loosely-associated materials, as in previous high throughput sequencing studies of *Cladophora* microbiota (Zulkifly et al. 2012, Graham et al. 2015a), algal filaments were washed three times in freshly made, autoclave-sterilized SD11, a defined medium closely resembling the chemical composition and pH of Lake Mendota water (Graham et al. 1996). The washing process involved using sterile forceps to transfer algal thalli into a sterile plastic petri dish containing sterile medium, swirling the dish for ∼1 minute, and repeating this process twice more, using fresh petri dishes and medium. *Cladophora* thalli were trimmed to retain only the most basal two cm, which included older basal cells and holdfasts: 1) to consistently sample host material of similar age, 2) because young host cells at growing tips are much less highly colonized, and 3) to facilitate comparisons with samples from two previous years, which had been similarly collected, processed, and analyzed (Zulkifly et al. 2012, Graham et al. 2015a). Cleaned, trimmed algal materials collected in 2014 were placed into 1.5 mL Eppendorf Safe-Lock tubes. We processed three technical replicates from each of the three microsites that were sampled once per week during two three-week periods, one period near the beginning and the other toward the end of the 2014 growth season (Julian days 172, 178, 185, 199, 206, and 214). Two of the 54 possible samples, one sample from site Point and one sample from site South on Julian day 206 (July 25^th^), were excluded from further processing due to contamination by soil from nearby shoreline, resulting in 52 final samples.

### DNA extraction, amplification, and sequencing

Total genomic DNA was extracted on the same day as collection and processing using the DNeasy Plant Mini Kit (Qiagen) amended with 100 µL of lysozyme (Sigma-Aldrich) of concentration 100 mg/ml, to increase access to DNA of bacterial cells having recalcitrant cell walls (Yuan et al. 2012). Samples included DNA from the host, epibionts DNA, and DNA of intracellular bacteria potentially present. All DNA was stored in a freezer at -20°C from the date of collection/extraction until September 16, 2014, when all DNA samples were transported to the University of Wisconsin-Madison Biotechnology Center for 16S rRNA gene amplification and amplicon sequencing. 16S rRNA gene is both highly conserved across the domain Bacteria and contains variable regions, making amplified reads of this gene ideal for conducting a broad taxonomic census of bacterial microbiota (Ward et al. 1990).

Variable regions V3 and V4 of the 16S rRNA gene were targeted as part of the 16S Metagenomic Sequencing Library Preparation Protocol, Par #15044223 Rev. B (Illumina Inc., San Diego, California, USA), using forward primer S-D-Bact-0341-b-S-17 and reverse primer S-D-Bact-0785-a-A-21 (Klindworth et al. 2013). Paired end, 250 base pair sequencing was performed using the Illumina MiSeq Sequencer and a MiSeq 500 bp (v2) sequencing cartridge. Amplicons were sequenced at the University of Wisconsin-Madison Biotechnology Center using the standard Illumina Pipeline, version 1.8.2, and these sequences have been deposited at the National Center for Biotechnology Information (NCBI) Short Reads Archive (SRA) under BioProject ID PRJNA360140.

### Bioinformatic processing

Forward and reverse paired ends were merged with PEAR (Zhang et al. 2014), and QIIME v.1.9.1 (Caporaso et al. 2010) was used to cluster reads with the UCLUST algorithm (Edgar 2010) and annotate resulting operational taxonomic units (OTUs) at 97% minimum similarity with the Silva database v. 128 (Quast et al. 2013). A total of 11,541,927 reads were obtained from amplicon sequencing. 985,143 reads were removed due to insufficient length or quality for assigning taxonomy. Taxonomy was assigned to the remaining 10,556,784 reads (a mean of 203,015 ± 47,471 reads per sample). Reads from Zulkifly et al. (2012), which was likewise a 16S rDNA amplicon study, were reanalyzed with the following bioinformatic processing for accurate comparison to the reads of the samples in this study.

Reads in each sample were filtered to represent only OTUs identified as bacterial in origin and singletons were removed. In an effort to reduce PCR bias that could inflate downstream OTU abundance metrics and indices, OTUs were normalized by relative abundance by dividing the sum of reads of each OTU by the sum of all reads in the sample. The Shannon index of alpha diversity, which incorporates both relative abundance and evenness across OTUs in a community (Sager & Hasler 1969), was calculated to measure site‑ and time-specific bacterial diversity. We chose the Shannon index for this study because the potentially low abundances of many rare OTUs detected from all bacteria associated with the alga influence the quantification of diversity (Wilhm 1970). The test for non-parametric multivariate analysis of variance (NPMANOVA) (Anderson 2001) was used to detect significant differences among microbial communities. Non-metric multi-dimensional scaling (NMDS) was used to cluster microbial communities, using a calculated Bray-Curtis dissimilarity distance matrix. All calculations and tests were completed using the R packages Phyloseq (McMurdie & Holmes 2013) and vegan (Dixon 2003).

### Comparisons of 2014 results to those reported from the same locale during 2011 and 2012

To gain insight into inter-annual variability, taxonomic results for samples collected in 2011 (Zulkifly et al. 2012) and 2014 (this study), which were obtained using similar amplicon methods, were compared at the class level. We also compared lists of bacterial genera reported from: 1) a 16S rRNA gene amplicon analysis of *Cladophora* microbiota sampled in 2011 (Zulkifly et al. 2012), 2) an analysis of samples collected from the same locale in 2012 using 16S rRNA gene sequences filtered from whole metagenomic shotgun sequence (Graham et al.2015a), and 3) the present analysis of *Cladophora,* sampled in 2014. This effort was designed to reveal possible core bacterial epibionts that may persist from year to year. However, in this process, the number of shared genera was almost certainly underestimated because public reference sequence databases grew during the 2011-2017 period encompassing the three contrasted studies.

We also employed the bacterial taxonomic data to infer the spatio-temporal dynamics of Cladophora-associated methanotrophic bacteria, whose activity in freshwater aquatic systems is of increasing interest. Because multiple methanotrophic bacterial genera-possibly reflecting functional redundancy‒were inferred for *Cladophora* microbiota sampled from the same site in 2011, yet a metagenomic analysis of *Cladophora* microbiota sampled in 2012 did not reveal this guild, we also examined the occurrence of methanotrophs in 2014.

## RESULTS

Epibiotic bacterial communities sampled from Lake Mendota (WI, USA) *Cladophora glomerata* at six time points and three microsites during the growing season of 2014 included 57,901 total possible OTUs present in 52 separate assemblages of 16S rRNA gene sequenced reads. The dominant bacterial phyla present in the *Cladophora* microflora, in decreasing rank by mean relative abundance, were *Bacteriodetes, Proteobacteria, Deinococcus-Thermus, Cyanobacteria, Acidobacteria,* and *Actinobacteria,* but *Bacteriodetes* and *Proteobacteria* accounted for most of the relative abundance (Supplementary Figure 3). Spatio-temporal occurrence of the 30 most abundant bacterial genera can be found in Supplementary Fig. 4.

In contrast to null hypotheses of spatial and temporal homogeneity, analysis of all reads identified as bacterial in origin indicated substantial temporal and spatial variation. The non‑parametric multivariate analysis of variance test showed that the bacterial microbiota associated with Picnic Point, Lake Mendota *Cladophora glomerata* differed significantly over the growth season (pseudo F = 26.05, R^2^ = 0.438, p < 0.001) and between microsites (pseudo F = 30.92, R^2^ = 0.208, p < 0.001). Because there was a significant interaction between sampling date and microsite (pseudo F = 7.11, R^2^ = 0.239, p < 0.001), Shannon diversity indices linear regressions were separated by microsite. The Point microsite showed a gradual net diversity increase over seasonal time (slope = 0.0144/day, R^2^ = 0.919, starting at 5.0, ending at 5.7). The South microsite more sharply increased in diversity over the growth season (slope = 0.0252/day, R^2^ = 0.9214, starting at 5.3, ending at 6.4) than did the North and Point microsites. The North microsite diversity decreased from 5.7 to 5.2 and ended at 5.6, but this trend did not fit a linear pattern (slope = 0.00263/day, R^2^ = 0.0851) (Fig. 1). 2014 samples clustered equally by collection date and site (Fig. 2), suggesting nearly equal contribution to differences in composition.

**Figure 1.**
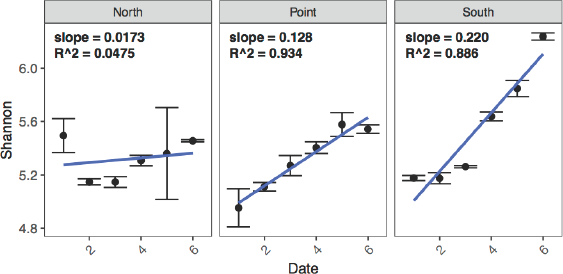
Mean alpha diversities, with linear regressions, as measured by the Shannon index of 16S amplicon operational taxonomic units (OTUs) that represent bacterial diversity calculated by weighting taxa richness by the evenness of the OTUs detected in the epibiota of *Cladophora* from three sites on Picnic Point, Lake Mendota, WI, during the summer of 2014. Error bars ±SE (n=3).

**Figure 2.**
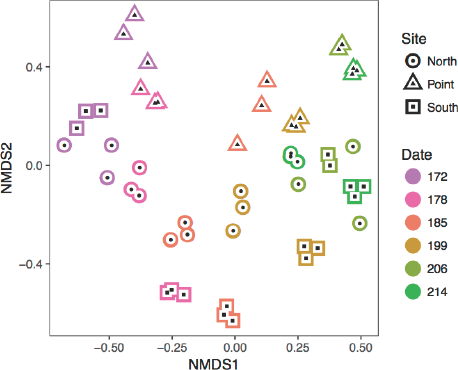
Non-metric multi-dimensional scaling (NMDS) ordination plot using Bray-Curtis distances of 16S rRNA gene amplicon reads identifed as bacterial, sequenced from samples of *Cladophora* collected from Picnic Point, Lake Mendota, WI, during the summer of 2014. Color indicates collection date (Julian days 172 to 214), and shape indicates collection site (circle = North, triangle = Point, square = South). This plot visualizes an ordination using Bray-Curtis calculated differences (k = 2) between all samples in two-dimensional space and has a sufficiently low stress of 0.1186.

However, in this study the collection period was relatively long, but distances between microsites were relatively small. The data indicate that samples from early-summer collection dates clustered separately from later-summer samples (Fig. 2).

At the class level, we observed considerable variability in relative bacterial abundance over time and space (Fig. 3). Notable trends of dominant classes were an abundance of *Alphaproteobacteria* at the Point microsite, and a general decline over time of *Acidomicrobia, Actinobacteria, Armatimonadia, Cytophagia,* and *Flavobacteria.* Re-analysis and comparison of class-level data originally obtained in 2011 (Zulkifly et al. 2012) suggested that abundances of taxa in classes *Acidomicrobiia, Actinobacteria, Clostridia, Deinococci, Deltaproteobacteria, Gammaproteobacteria, Gemmatimonadetes, Nitrospira,* and *Sphingobacteria* were lower in 2011 than nearly all sampled dates in 2014 (Fig. 3). The same contrast indicated that abundances of classes *Anaerolineae, Betaproteobacteria,* and *Flavobacteria* were all higher in 2011 than at any time in 2014 (Fig. 3).

**Figure 3.**
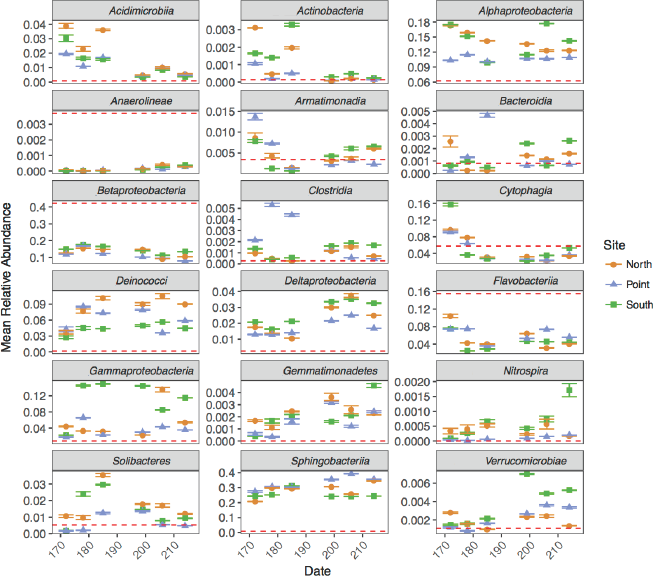
Mean relative abundance of epibiont bacterial classes on *Cladophora* sampled from Picnic Point, Lake Mendota, WI, during the summer of 2014, inferred from 16S rRNA gene sequences. Error bars ±SE (n=3). Red dashed lines indicate values for classes detected by 16S rRNA gene sequences associated with *Cladophora* collected from the same locale in the summer of 2011 by Zulkifly et al. (2012) after reanalysis using the bioinformatic processing used in this study.

A total of 33 consistently detected bacterial genera associated with *Cladophora* were found in 2011 (Zulkifly et al. 2012), 2012 (Graham et al. 2015), and 2014 (this study), and these genera are listed in Supplementary Table 1. Seven putatively methanotrophic genera were inferred to occur as epibionts on *Cladophora* sampled in 2014, listed here in order of greatest to least mean relative abundance: *Crenothrix, Methylobacter, Methylovulum, Methylocaldum, Methylocella, Methylocystis,* and *Methylomonas.* Relative abundances of these genera varied through the growth season and across microsites. Methanotrophic bacterial genera were inferred to be most abundant at the South microsite throughout the summer of 2014 and in early summer at the North microsite, but were nearly undetectable at the Point microsite (Fig. 4).

**Figure 4.**
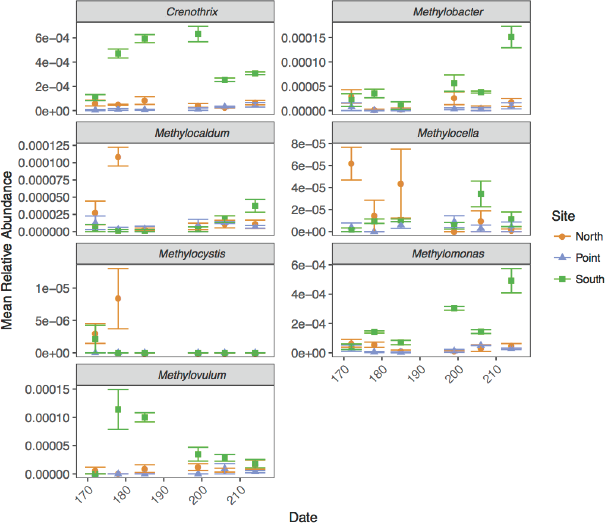
Mean relative abundance of methanotrophic (methane-oxidizing) genera occurring in the bacterial microbiota of *Cladophora* sampled at three microsites (North, Point, and South) from the locale Picnic Point, Lake Mendota, WI, during the summer of 2014, inferred from the 16S rRNA gene. Error bars ±SE (n=3).

## DISCUSSION

The goals of this study were to statistically test the null hypotheses that lacustrine *Cladophora* glomerata-associated bacterial communities were relatively homogeneous across the growing season of 2014, and that communities were relatively homogenous among relatively close microsites at the same sampling locale. Published taxonomic data from two previous years were also used together with 2014 information to gain insight into inter-annual variation, with a particular focus on methanotrophs, because methane-oxidation is known to be a key ecosystem function in lakes (Khmelenina et al. 2010, Deutzmann et al. 2014, Billard et al. 2015, Zigah et al. 2015, Yang et al. 2016, Roland et al. 2017, Samad and Bertilsson 2017).

### Temporal and spatial variation

Our results indicated that populations of relatively shallow lacustrine *Cladophora* support a highly dynamic bacterial community whose composition within a growing season was influenced twice as much by sampling day as by closely-placed microsites. The results indicated that future studies of periphytic algal microbiota should include more than two technical replicates per sample site as well as sampling at multiple times during a growth season. Variation also occurred among epibiotic bacterial communities of closely-spaced microsites, but optimal spacing of sampling sites may depend upon the specific hypotheses being tested and upon the degree of variation in locale features such as substratum quality and shading, and therefore may need to be determined for individual ecosystems.

The consistent increase in alpha diversity we observed across two microsites during the course of a growth season suggests a pattern of accumulation of bacterial taxa. Even so, observed increases and decreases in relative abundance at all taxonomic levels suggests that complex successions of microbial guilds may surge and retreat in response to microenvironment changes. Environmental factors other than water temperature were not assessed in our study, but future investigations of periphytic algal microbiota might also include monitoring of variation in sunlight, wave action, shoreline aspect, human use, and shoreline runoff, to begin to parse the drivers of community change over time.

### Functional inferences

Although there are limitations to inferring bacterial functions from taxonomic information (Langille et al. 2013), inferring function from generic classification can be justified when a metabolic function is highly conserved at the generic level. Our observations of bacterial communities associated with lacustrine *Cladophora* during the growing season of 2014 reveals patterns of change in bacterial genera that might be ecologically significant and thus warrant further investigation. For example, the studied eutrophic lake is rich in dissolved organic carbon, phosphorus, and nitrogen (Beversdorf et al. 2013; Carpenter et al. 2014; Karatayev et al. 2013), which may foster the occurrence of hardy copiotrophs such as *Meiothermus* (Tindall et al. 2010), *Sediminibacterium* (Kang et al. 2014), *Pseudomonas* (Klausen et al. 2003), and *Flectobacillus* (Hwang et al. 2006), all of which occurred at high relative abundance on *Cladophora.* The phototrophic cyanobacterium *Leptolyngbya* (Shimura et al. 2015) occurred in greater relative abundance at the least-shaded microsite, but *Rhodobacter,* commonly a non-oxygenic photosynthesizer (Hiraishi et al. 1996), did not seem to have a preferred microsite.

Temporal and spatial patterns of methanotrophs may be of particular functional importance. Although most studies of lacustrine methanotrophs have focused on suspended or benthic taxa (Khmelenina et al. 2010, Deutzmann et al. 2014, Billard et al. 2015, Zigah et al. 2015, Yang et al. 2016, Roland et al. 2017, Samad and Bertilsson 2017), our past (Zulkifly et al. 2012) and present evidence for multiple genera of putatively methanotrophic bacteria in *Cladophora* epibiotic communities indicates that littoral methanotrophs may also be important. Our inferences of rapid increase followed by slow decrease in the relative abundances of the putative methanotrophs *Crenothrix* (Stoecker et al. 2006), *Methylocaldum* (Islam et al. 2015), and *Methylomonas* (Ogiso et al. 2012) suggest that littoral methane concentrations may have changed over time, possibly reflecting differences in emissions from sediments. The putative methanotrophs *Methylomicrobium* (Brantner et al. 2002) and *Methylosinus* (Fox et al. 1989) were found in lower relative abundance, but these genera contributed to distinctive community responses at the South microsite. These results indicate that greater attention should be paid to methanotrophs associated with periphytic algae, particularly for lakes having a high proportion of littoral zone occupied by *Cladophora* and other periphytic algal hosts.

### Inter-annual comparisons

Consistent with our two previous, less well-replicated studies that were conducted at the same locale (Zulkifly et al. 2012, Graham et al. 2015a), the present assessment indicated that the microbiota of *Cladophora glomerata* includes representatives of diverse bacterial phyla, but that *Bacteroidetes* and *Proteobacteria* dominated. Although we inferred presence of similar bacterial classes during the 2011 and 2014 seasons, differences in mean relative abundance were observed in the two years, suggesting inter-annual variation. However, because our results indicated that bacterial communities can vary even among closely‑located microsites sampled on the same day, slight differences in collection location employed in the different years might explain some of this variation. In addition, less well-replicated sampling in the earlier year might have resulted in time‑ or site-biased collection bias. Even so, future explorations of inter-annual variation in *Cladophora* epibiotic bacterial communities and possible environmental correlates seem justified in view of potentially powerful ecosystem functions such as methanotrophy.

Our observation that >30 bacterial genera, including functionally important representatives (Zulkifly et al. 2012), were present on *Cladophora* in all three study years suggests the occurrence of a core microbiota that might be transmitted from one year to the next. The mechanism of transmission might be persistent thick-walled akinetes that remain attached to submergent substrata from year to year (Zulkifly et al. 2013). To test this hypothesis, future investigations might explore the microbiota of *Cladophora* akinetes collected during late fall, winter, and early spring, to check for the ∼30 putative core bacterial genera.

The contrast between inferred presence of methanotrophic bacteria in epibiotic *Cladophora* communities in 2011 and 2014 and complete lack of detection of methanotrophs or sequences encoding enzymes that mark methane oxidation (e.g. *pMMO)* in similarly collected and processed samples from 2012 suggests an important environmental difference between these years. Graham et al. (2015a) suggested that an earlier than usual rise in lake temperature coupled with a longer than usual period of sustained maximal lake temperature that occurred in 2012 may have been detrimental to the Cladophora-methanotroph association. However, because the present study indicated that putative methanotrophs can be abundant at one microsite, yet undetectable at a nearby microsite, it is possible that 2012 spatial sampling missed methanotrophs that might have been present elsewhere. These observations suggest that future efforts to monitor the abundances, activities, and spatial and temporal heterogeneity of Cladophora-associated methanotrophs are warranted. Even though conspicuous *Cladophora* growths in freshwater bodies are often maligned, it is becoming increasingly more evident that the holobiont serves important ecosystem functions such as methanotrophy.

## ACKNOWLEDGEMENTS

We thank the Anna Grant Birge Fund administered by the UW-Madison Center for Limnology financial support for amplicon sequencing, and UW-Madison Biotechnology Center personnel for services and expertise in Next-Gen Sequencing technology.

## Supplementary Information

**Supplementary Figure 1.**
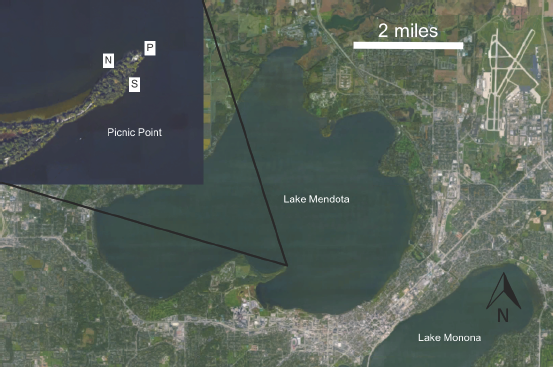
Map of collection sites North (N), Point (P), and South (S) on Lake Mendota, WI, USA.

**Supplementary Figure 2.**
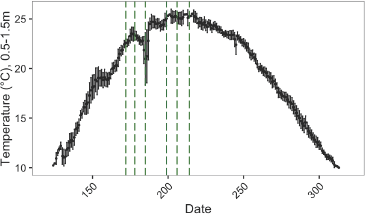
Mean daily temperatures aggregated from the Mendota Buoy Long Term Ecological Network database from 2006 to 2016, excluding data from 2014 when intrumentation malfunctioned, resulting in large temperature measurement gaps and errors. Green dashed lines indicate sampling dates of this study.

**Supplementary Figure 3.**
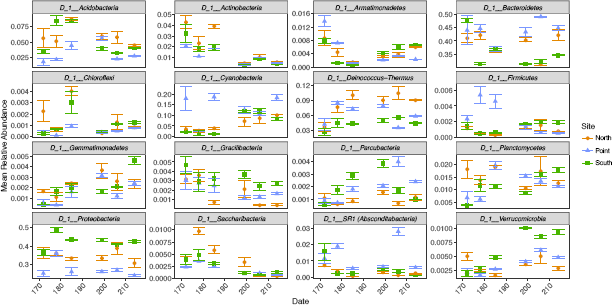
Mean relative abundances of the 16 most abundant bacterial phyla occurring in the bacterial microbiota of *Cladophora* sampled at three microsites (North, Point, and South) from the locale Picnic Point, Lake Mendota, WI, during the summer of 2014, inferred from the 16S rRNA gene. Error bars ±SE (n=3).

**Supplementary Figure 4.**
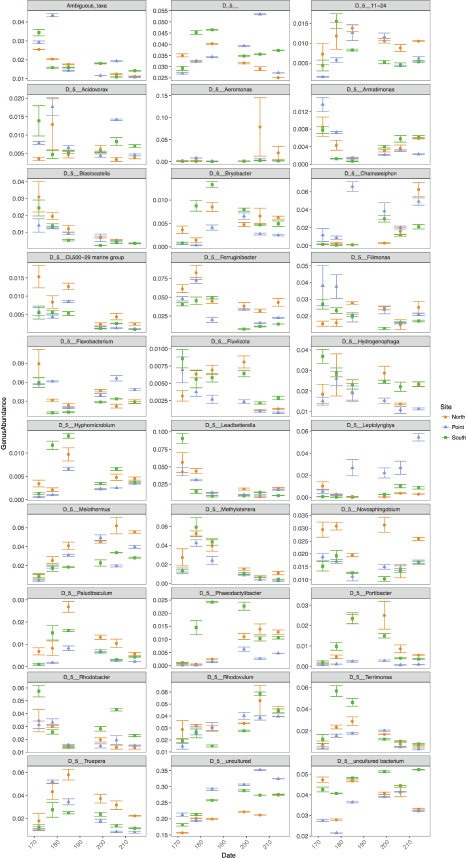
Mean relative abundances of the 30 most abundant bacterial genera occurring in the bacterial microbiota of *Cladophora* sampled at three microsites (North, Point, and South) from the locale Picnic Point, Lake Mendota, WI, during the summer of 2014, inferred from the 16S rRNA gene gene. Error bars ±SE (n=3).

**Supplementary Table 1.**
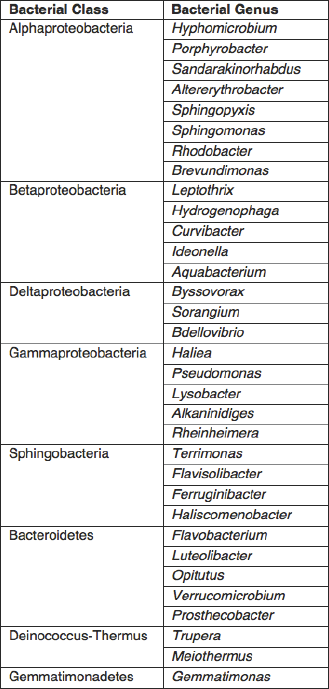
Recurring core bacterial genera closely associated with *Cladophora glomerata* in Lake Mendota identified using short reads of the 16S rRNA in the growth seasons of 2011 (Zulkifly et al. 2012), 2012 (Graham et al. 2015), and 2014 (this study).

